# MOH: a novel multilayer multi-omics heterogeneous graph for single-cell clustering

**DOI:** 10.1101/2025.08.04.668248

**Authors:** Yao Dong, Chen Chen, Jin Shi, Yushan Hu, Xiaowen Cao, Yongfeng Dong, Xuekui Zhang

## Abstract

Cell clustering is crucial in single-cell multi-omics research for identifying distinct cellular populations. Although there has been progress in integrating multi-omics data for clustering, combining more than two types of omics data remains challenging due to the diversity and heterogeneity of these datasets. Traditional approaches typically use heterogeneous graphs that integrate only two types of omics data, constructing graphs with genes and cells as nodes and a single type of edge representing their relationships. However, this method has limitations as it overlooks cell-cell interactions and struggles to capture complex cellular dynamics. Additionally, the graph structure must be redesigned whenever new omics data are introduced, limiting the scalability of these models. To address these issues, we introduce MOH, a novel single-cell clustering algorithm based on a multilayer multi-omics heterogeneous graph. MOH integrates three key single-cell omics types: scRNA-seq, scATAC-seq, and spatial transcriptomics. It constructs a multilayer heterogeneous graph to simultaneously extract and enhance representations from all three omics layers, incorporating both intra-layer and inter-layer edges to capture association and similarity relationships. This enriched representation leads to an accurate clustering results. Extensive experiments show that MOH outperforms six state-of-the-art methods on unsupervised clustering metrics, offering a precise and comprehensive analysis with consistent improvements across all evaluation criteria. Moreover, downstream analyses validate the results, revealing novel biological insights into immune disorder complications in cancer, cancer drug repurposing, and new signaling pathways, which merit further investigation and validation.

## I. Introduction

**T**he advancement of single-cell genomics technology offers extremely detailed insights at the cellular level. An important initial task in analyzing single-cell data is accurately identifying the type of each cell, as subsequent analyses may lack context without precise cell type annotation [1]. Therefore, the accuracy and reliability of all subsequent analyses in single-cell studies depend on correctly identifying the cell type in the initial step [2]. Clustering is a widely used method for annotating cell types in the analysis of single-cell genomic data. This unsupervised method is crucial for identifying new cell populations, understanding cellular heterogeneity, and discovering rare cell types within complex tissues.

While traditional genomics or transcriptomics studies offer valuable insights, they often capture only a single dimension of cellular activity. The popularity of multi-omics data is rapidly increasing due to its ability to provide a more comprehensive and nuanced view of biological systems. Multi-omics approaches, which can include transcriptomic (scRNA-seq) [2], spatial transcriptomic (ST) [3], epigenomics (e.g. scATAC-seq) [4], etc., enable the simultaneous analysis of multiple biological layers. Integrating multi-omics leads to more precise and biologically meaningful clustering [5], enhancing our understanding of cellular diversity and function [6].

However, significant challenges remain in combining these diverse and heterogeneous datasets [7], including data format heterogeneity, technical noise, batch effects, and computational scalability [8]–[10]. Effective multi-omics integration is essential for advancing single-cell clustering techniques, yet current methods predominantly focus on integrating two of these three omics layers, including:

a. Integrating scRNA-seq and scATAC-seq: This approach collectively depicts the transcriptional regulatory landscape of cells, using shared cell labels [11] to align cellular identities across modalities, gene activity scores [12] to convert chromatin accessibility into transcriptome-compatible features, and graph-based alignment techniques [13] to preserve high-dimensional relationships between omics. However, these methods still face challenges in handling cellular heterogeneity and data sparsity, especially in complex biological systems. Moreover, the lack of spatial information prevents the revelation of gene regulation and expression distribution within tissue spatial structures. To address the above issues, spatial transcriptomics technologies, including high-resolution ST and multi-modal imaging, are advanced. High-resolution ST have significantly improved spatial gene expression localization, addressing limitations of earlier spot-based techniques that mixed signals from multiple cells. Multi-modal imaging now enables coregistration of spatial transcriptomics with proteomics and chromatin accessibility, providing a more complete view of cellular microenvironments. The two typical multi-modals with ST are as follows:
b. Integrating scRNA-seq and ST data: Integrating scRNA-seq with ST data was challenging due to differences in resolution and the complexity of aligning datasets. The latest development stage emphasizes improving resolution and utilizing graph-based methods for spatial data analysis. Graph-based spatial integration techniques enable better modeling of cell-cell interactions and address challenges in spatial data alignment. Tools like SpatialDE [16] and Seurat [17] now enable joint analysis of scRNA-seq and ST data, leveraging high-resolution ST to map single-cell gene expression profiles onto spatial contexts. Nevertheless, the absence of chromatin state information in these integrations limits understanding of upstream regulatory mechanisms and chromatin accessibility.
c. Integrating scATAC-seq and ST data: Traditional ST methods provided only gene expression data, limiting the ability to study the interplay between gene expression, protein localization, and chromatin accessibility. This approach aims to correlate the spatial information of chromatin accessibility with gene expression, deepening the understanding of the spatial dimension of gene regulation. Current methods, such as SnapATAC [19] and SpatialATAC [20], utilize joint analysis frameworks and multimodal graph models to identify spatial commonalities across omics datasets, attempting to integrate scATAC-seq and ST data. However, this integration lacks specific gene expression information, making it difficult to fully understand cells’ functional states and molecular characteristics.

Current methods handling more than 2 omics simply concatenate data matrix of different omics, which has two key disadvantages: (1) only incorporate “shared” information potentially masking important “omics-specific” biological signals [7], [21]. (2) concatenating generate to higher dimensional pooled data leads to overfitting [22], [23].

To address these challenges, we propose a novel <u>M</u>ultilayer multi-<u>O</u>mics <u>H</u>eterogeneous graph for single-cell clustering incorporating scRNA-seq, scATAC-seq, and ST data, named MOH. Inspired by heterogeneous graph learning, we construct a multilayer heterogeneous graph that connects three types of omics nodes and extracts intra-omics and interomics associations and similarities. A multilayer heterogeneous graph enables structured integration of multiple omics while preserving modality-specific information and enabling cross-omics interaction, improving clustering accuracy. The major contributions of this work are summarized below:

1. The capability to extend to additional omics types: We integrate more than two omics data, including scRNA-seq, scATAC-seq, and ST data, with the capability to extend to additional omics types. Our approach employs advanced integration techniques including cross-layer cell information sharing through spatial and molecular similarity alignment, MAESTRO-derived regulatory potential weights for transforming chromatin accessibility to gene activity scores, and Weighted Nearest Neighbors (WNN) methodology for balanced multi-modal integration. These techniques significantly improve clustering performance while effectively preserving the unique biological information inherent to each omics type.
2. Construction of a multilayer heterogeneous graph: We introduce a novel multi-layer graph architecture that organizes different omics data into distinct but interconnected layers. The first layer integrates scRNA-seq and scATAC-seq data, while the second layer represents ST data. Our design uniquely features cross-layer cell mapping and inter-layer edges, creating a unified representation while maintaining modality-specific characteristics. Importantly, this architecture is inherently extensible, allowing seamless integration of additional omics layers.
3. Enhanced representation in heterogeneous graphs: We enhance the representational content of heterogeneous graphs by introducing additional edges. Traditionally, heterogeneous graphs used to integrate scRNA-seq and scATAC-seq data include only one type of edge, namely cell-gene edges. To improve the learning of intra-layer interactions, we introduce new cell-cell edges within each layer. This enhancement strengthens intra-layer connectivity, enriching cell representations and providing a more comprehensive understanding of intracellular dynamics and interactions.
4. Novel biological results generated by downstream analysis of MOH: Based on the results of MOH, we conduct downstream analyses to validate the correctness of MOH’s cell type annotation and demonstrate how to generate new knowledge. Specifically, we generated novel hypotheses in complications of immune disorder in cancer, cancer drug repurposing, and new signalling pathways, which can be investigated and validated in further studies.

## II. Related Work

Each type of omics data in single-cell multi-omics possesses unique characteristics, and constructing data graphs can reveal multifaceted biological information from different perspectives. Researchers build biological networks from various omics datasets to enhance clustering and improve system-level understanding. Recently, the biological network inference model Deepmaps [27], based on heterogeneous graphs and heterogeneous graph transformers, and the graph convolutional network model CCST [28], which learns spatial structures, has demonstrated superior performance.

The Deepmaps model employs heterogeneous graphs and heterogeneous graph transformers to infer biological networks from multimodal omics data, utilizing multi-head graph transformers to robustly learn the relationships between cells and genes in both local and global contexts. CCST is tailored for ST data and is a cell clustering method based on GCNs. It integrates single-cell gene expression with intricate global spatial information from spatial gene expression data.

Despite their strengths, both Deepmaps and CCST have limitations. Deepmaps does not incorporate ST data, thus missing out on learning global spatial information. Conversely, CCST focuses exclusively on ST data, limiting the diversity of input omics data. To address these shortcomings, we propose a novel multilayer multi-omics heterogeneous graph for single-cell clustering, named MOH. Our approach synergizes the strengths of Deepmaps and CCST by integrating three types of omics data. MOH constructs a multilayer heterogeneous graph, expanding the fusion of the three omics datasets and learning the specific information from each omics data type.

## III. Methods

MOH employs heterogeneous graphs to represent and analyze multi-omics data at the single-cell level. The heterogeneous graph contains different types of nodes and edges, allowing it to capture diverse biological relationships in the data. The architecture of MOH is shown in Fig. 1. It consists of three key modules: the scRNA-seq and scATAC-seq heterogeneous graph module, the ST graph module, and the multi-omics multilayer heterogeneous graph for cell clustering module. The first two modules work synergistically to process different data modalities, including scRNA-seq and scATAC-seq data, and ST data respectively. The last module integrate them into a unified representation, ultimately enabling comprehensive cell state characterization. The following sections detail the specific methodologies of each module.

**Fig. 1:**
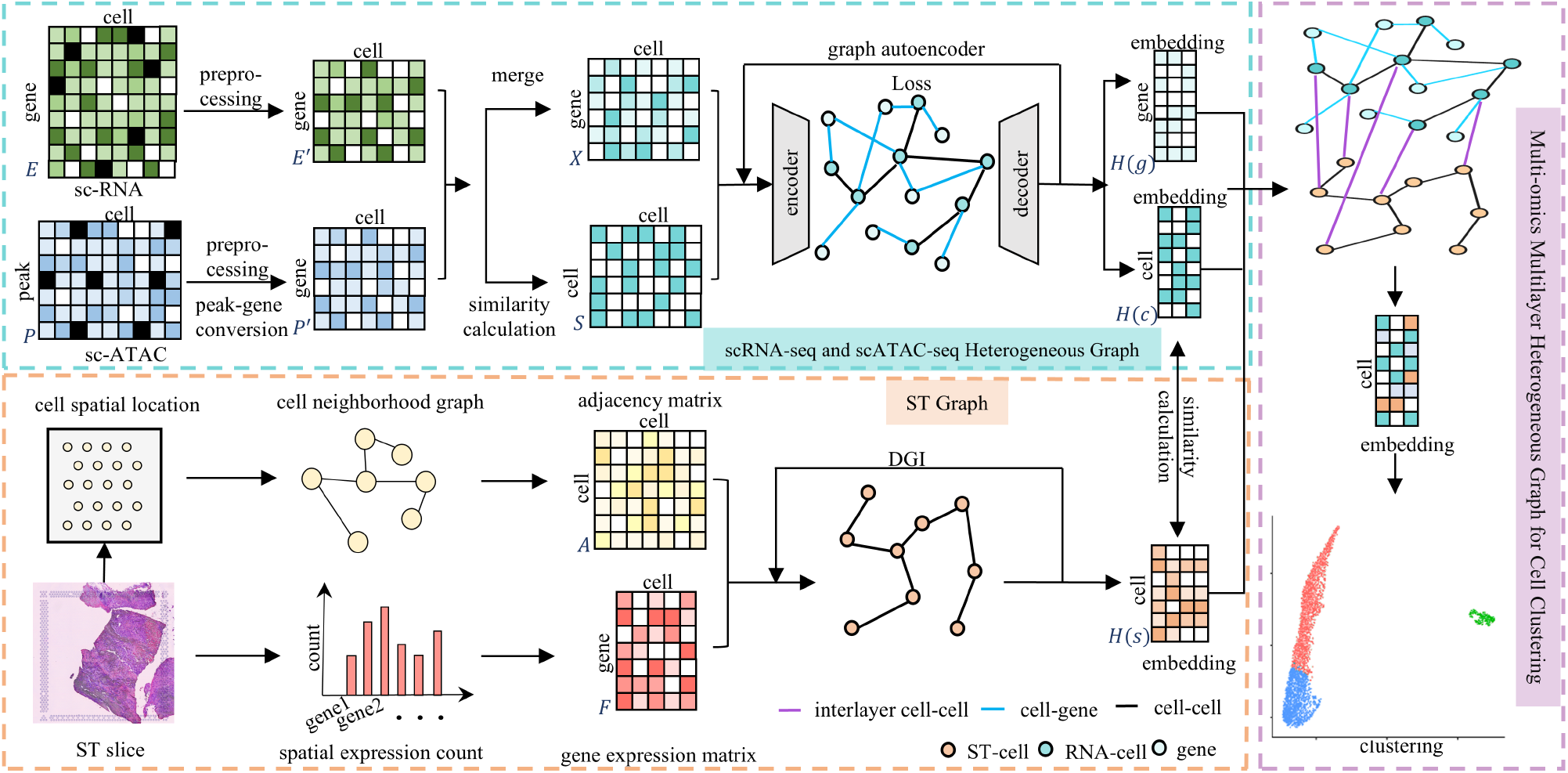
Architecture of MOH. The scRNA matrix *E* and scATAC matrix *P* are preprocessed and integrated together through peak and gene weight transformations to construct a heterogeneous graph. The graph contains two types of nodes, cells and genes, and connected by two edges, cell-cell and cell-gene. GAE uses this graph to obtain cell embeddings *H*(*c*) and gene embeddings *H*(*g*) through communication between nodes within the layer. The ST graph construction module calculates the cell position and gene count data of spatial cell slices to construct the graph, and obtains the cell embedding *H*(*s*) of ST through GCN. Subsequently, a multilayer heterogeneous graph is constructed by mapping cells between layers. Node embeddings are enhanced through inter-layer learning to obtain the final embedding for clustering.

### A. scRNA-seq and scATAC-seq Heterogeneous Graph

We construct the scRNA-seq and scATAC-seq heterogeneous graph using 10X platform’s single-cell multiome ATAC+ gene expression data. The cells in the two types of omics data are matched. This input includes the raw matrices: scRNA-seq (gene expression) and scATAC-seq (chromatin accessibility).

#### 1) Data Processing

For preprocessing, we utilize Seurat v4 only for basic data preprocessing steps including normalization, scaling, and dimensionality reduction. We convert the original scRNA-seq matric *E* and original scATAC-seq matric *P* into cell-gene matrices *E*^*′*^ and *P* ^*′*^ with the same dimensions. Each data matrix consists of cells (columns) and genes or peaks (rows). Rows or columns with fewer than 0.1% non-zero values are removed. Gene expression matrices are log-normalized, and the top 2000 highly variable genes are identified using Seurat v4 for each matrix. If a matrix contains fewer than 2000 genes, all available genes are selected for integration [27].

For scRNA-seq, we denote the gene expression matrix as *E* = {*e*_*ij*_ |*i* = 1, 2, …, *I*; *j* = 1, 2, …, *J*}, where *I* and *J* represent the number of genes and cells, respectively. To obtain a qualitative representation of each gene across all cells, we employ a Hidden Markov Model (HMM). In this model, the expression state of each gene *i* can be discretized into *M*_*i*_ states. Specifically, the HMM defines and generates *M*_*i*_ discrete states, representing different regulatory signals or expression levels of gene *i* across all *J* cells. By applying the HMM, we can produce *M*_*i*_ distinct states for each gene, capturing its discretized expression patterns across different cells.

For scATAC-seq data, we name the chromatin accessibility matrix *P* = {*p*_*kj*_ |*k* = 1, 2, …, *K*; *j* = 1, 2, …, *J*}, *k* and *j* are used to denote the peaks and cells respectively. To integrate scRNA-seq and scATAC-seq data, we aim to transform *P* into a gene-cell matrix *P* ^*′*^. Firstly, we employ the MAESTRO [29] method to calculate the regulatory potential weight *W*_*ki*_ from peak *k* to gene *i*:

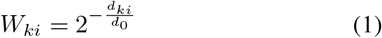

where *d*_*ki*_ is the distance from the center of peak *k* to the transcription start site (TSS) of gene *i*, and *d*_0_ is a user-defined half-decay distance, typically set to 10 kb. For a given gene *i*, if the distance *d*_*ki*_ from peak exceeds 150 kb, the weight *W*_*ki*_ is set to less than 0.0005 when *d*_0_ is 10 kb. In our method, to optimize computational efficiency, *W*_*ki*_ is set to 0 if the distance between the peak and TSS exceeds 150kb [27].

For peaks located within exonic regions, we adjust the weights to account for gene length biases. If peak *k* is in the exonic region of gene *i*, its weight is normalized by the total exon length to correct for the higher probability of peak detection in longer genes. Conversely, peaks in promoter or exonic regions of nearby genes are excluded from contributing to the gene’s regulatory potential.

To accurately reflect the regulatory potential, we adjust for biases introduced by the differing lengths of gene regions by normalizing the weight of each peak. This normalization accounts for the varying detection probabilities across different gene regions and provides a more precise measure of regulatory activity. The final regulatory potential matrix *P* ^*′*^ is then computed by combining these adjusted weights with the original peak matrix to obtain gene-level regulatory scores.

#### 2) Data Integration

We employ the Weighted Nearest Neighbors (WNN) [34] approach to integrate the sc-RNA and sc-ATAC matrices. The WNN integration method leverages the principal components from both omics datasets to construct a shared nearest neighbor graph. Let the gene expression matrix in the PCA-reduced space be 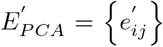, where *i* = 1, 2, …, *I* represents the principal components, and *j* = 1, 2, …, *J* represents the cells. For each cell *j*, the Euclidean distance 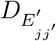 to every other cell *j*^*′*^ is calculated as follows:

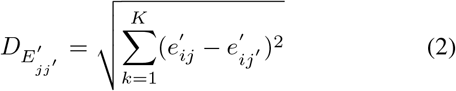

The same action is taken for 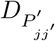.

For each cell *j*, the weights 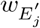 is calculated as follows:

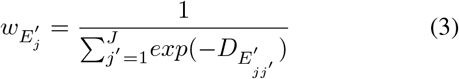

The same formula applies to 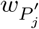. These weights are normalized to ensure that the sum of weights for each cell equals 1.

Each edge in the WNN graph is weighted by the respective distances from the gene expression and chromatin accessibility KNN graphs. The combined distance 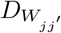 for cell *j* is calculated as a weighted sum of the individual distances:

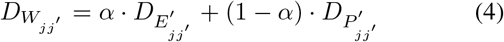

where *α* is a parameter that balances the contributions from both data types, we set *α* = 0.5 [11].

Using the combined distances *D*_*W*_, we construct a binary nearest neighbor matrix *N*_*W*_, where each entry indicates whether two cells are nearest neighbors (1) or not (0) based on their weighted combination of gene expression and regulatory potential distances.

To integrate the gene expression and chromatin accessibility matrices into a single gene-cell matrix, we leverage the WNN graph. For each cell *j*, its integrated expression profile *x*_*j*_ is obtained by averaging the profiles of its nearest neighbors from both datasets:

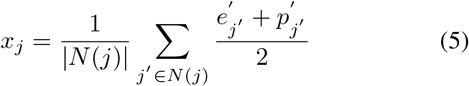

where *N* (*j*) represents the set of nearest neighbors of cell *j* in the WNN graph, 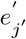 is the gene expression vector of cell *j*^*′*^ from sc-RNA data, and 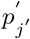 is the gene expression vector of cell *j*^*′*^ from sc-ATAC data.

To simplify notations, we redefine the integrative matrix generated in the previous section as *X* = {*x*_*ij*_ | *i* = 1, 2, …, *I*; *j* = 1, 2,, …, *J*}, where *i* denotes the number of genes and *j* denotes the number of cells. To understand the relationships between cells, we compute the cosine similarity of their expression features. This involves calculating pairwise cosine similarities between cells based on their gene expression vectors. The cosine similarity 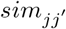 between cell *j* and cell *j*^*′*^ is defined as:

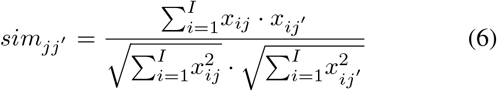

Using these distances, we construct a cell-cell similarity matrix *S*. where each entry is set to 1 if the similarity between two cells is non-negative, and 0 otherwise.

#### 3) Graph Construction

We construct a heterogeneous graph *G* using the gene expression matrix *X* and the similarity matrix *S*. Nodes in graph *G* represent cells and genes, with edges depicting cell-cell similarities and cell-gene expression relationships. We use the gene expression vectors of cells as the initial embedding 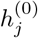 for cell nodes and the expression levels of genes across all cells as the initial embedding 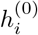 for gene nodes. Initial embeddings 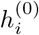 initialize the Graph Autoencoder(GAE). The role of the initial embeddings is to provide the encoder with meaningful features derived directly from the biological data, ensuring that the initial representation of each node captures important information about its gene expression profile.

We employ a GAE to extract meaningful low-dimensional representations. The GAE comprises an encoder and decoder. The encoder maps nodes in the graph *G* to a low-dimensional space. For each node *v* (which can be either a cell or a gene), the encoder generates a low-dimensional embedding *H*(*v*), formally defined as 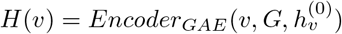.

The GAE is trained by minimizing the reconstruction loss, which measures the decoder’s ability to reconstruct the original graph structure from the embeddings generated by the encoder. The loss function *L* is defined as:

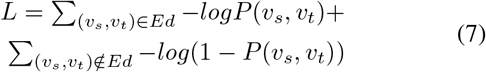

where *Ed* is the set of edges in the original graph, and (*v*_*s*_, *v*_*t*_) represents a pair of nodes.

After completing the training, the encoder provides the embeddings for cells *H*(*c*) and the embeddings for genes *H*(*g*). Then, these low-dimensional embeddings can be utilized for various downstream tasks, such as clustering, visualization, or further biological analysis.

### B. ST Graph

The input for the spatial transcriptomics graph construction module is ST (spatial transcriptomics) data generated by the 10X platform’s product, Spatial Gene Expression. We preprocessed the input ST data by filtering out low-expressed and low-variance genes and normalizing the counts for each cell. The ST graph can be represented by two matrices: the cell location adjacency matrix and the node feature matrix.

First, we calculate the distance *d*_*ij*_ between each pair of cells (cell *i* and cell *j*), and select an appropriate threshold *d*_0_ to construct the cell location adjacency matrix *A* = {*a*_*ij*_ |*i* = 1, 2, …, *N* ; *j* = 1, 2, …, *N*}, where *N* is the number of cells. The matrix entries are binary, with 1 indicating that the distance between two cells is below a selected threshold *d*_0_, and 0 otherwise.

The cell location adjacency matrix constitutes an undirected graph that represents the spatial information between cells, where cells are nodes and spatially adjacent cells are connected by edges.

The node feature matrix *F* = {*f*_*ij*_ |*i* = 1, 2, …, *I*; *j* = 1, 2, …, *N*} is derived from the preprocessed gene count data, which is then standardized:

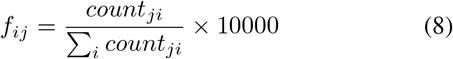

where *i* represents gene*i, j* represents gene*j*. Then, genes with a standardized expression variance lower than 1 are removed to exclude low-variance genes.

We use Deep Graph Infomax (DGI) [35], an unsupervised graph embedding method, to learn node representations from graph-structured data. Unlike GCN-based methods that rely on random walks, DGI maximizes the mutual information between global and local representations of the graph. The features extracted by DGI include both local and global.

The combined adjacency matrix *A* and the node feature matrix *F* are used as inputs to DGI. The encoder maps these two matrices to an embedding space:

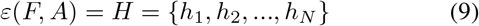

where *H* represents high-order representation.

The encoder is composed of four graph convolutional layers for passing messages over neighboring nodes with a Parametric Rectified Linear Unit (PReLU) as the activation function. The graph convolutional layer is defined as follows:

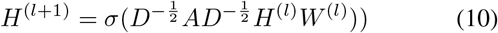

where *H*^(*l*)^ and *H*^(*l*+1)^ are the input and output matrices, *W* ^(*l*)^ is the weight matrix, and *D* is the degree matrix.

The PReLU activation function is:

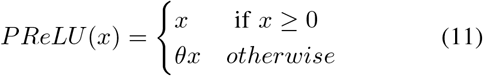

where *θ* is a learnable parameter, we set *θ* = 0.25 [27].

Global representation is derived from local representations using a readout function:

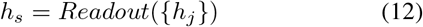

A discriminator is introduced to maximize the mutual information between local and global features, with higher scores indicating better encapsulation of graph-level information. To train the discriminator, negative samples Ã are generated by randomly corrupting the adjacency matrix *A*. The objective function to be maximized is:

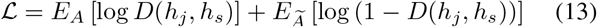

where *D* is the discriminator function. Thus, we obtain the output matrix *H*(*s*), which is the representation matrix of cells.

### C. Multi-omics Multilayer Heterogeneous Graph for Cell Clustering

Let the output cell embedding matrix from Module A (scRNA-seq and scATAC-seq Heterogeneous Graph Construction) be defined as *H*(*c*) = {*h*(*c*)_*k*_| *k* = 1, 2, …, *K*, and the gene embedding matrix be defined as *H*(*g*) = {*h*(*g*)_*i*_ | *i* = 1, 2, …, *I*}, where *K* is the number of cells input into Module A and *I* is the number of genes. The gene embedding matrix *H*(*g*) extracted from Module A plays an essential role in enriching the node representations by providing additional gene-level information during the integration process. Similarly, let the output cell embedding matrix from Module B (Spatial Transcriptomics Graph Construction) be defined as *H*(*s*) = {*h*(*s*)_*j*_| *j* = 1, 2, …, *J*}, where *J* is the number of cells input into Module B.

To facilitate the integration of these matrices, we first reduce both matrices *H*(*c*) and *H*(*s*) to the same dimensionality using Principal Component Analysis (PCA). Specifically, we reduce them to 128 dimensions: *H*(*c*) and *H*(*s*) have dimensions *K ×* 128 and *J ×* 128, respectively.

Next, we calculate the similarity between the cells in the reduced matrices *H*(*c*) and *H*(*s*) using cosine similarity. For the *k*-th cell in *H*(*c*) and the *j*-th cell in *H*(*s*), the cosine similarity *s*_*kj*_ is given by:

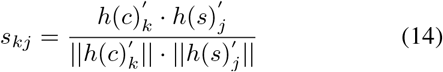

To fully leverage both inter-layer cell similarities and gene-level information, we define update rules that incorporate both the cross-layer cell embedding similarities and gene embeddings *H*(*g*). The gene embeddings provide crucial biological context that helps refine the cell representations. For each cell node *k* in matrix *H*(*c*), its embedding is updated based on both cross-layer similarities and gene-level information as follows:

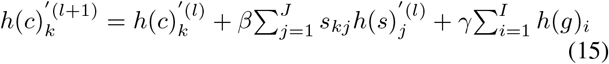

Similarly, we can obtain the update rule for cell node k in matrix *H*(*s*) following the same approach, where *β* and *γ* are learning rates that control the contribution of cross-layer similarity and gene embeddings, respectively. In experiment, we added comprehensive parameter experiments for *β*. The results demonstrate optimal performance at *β* = 0.4 across all evaluation metrics. For *γ*, we implemented a data-driven dynamic adjustment strategy.

This iterative update continues until convergence, ensuring that the cell embeddings in both layers are mutually informed by their cross-layer connections and gene information, thereby refining the heterogeneous multi-omics graph. The key advantage of our approach lies in its ability to simultaneously preserve layer-specific information while enabling effective information sharing across layers. This balance is achieved through the careful design of update rules that consider both cross-layer similarities and gene-level context, resulting in more comprehensive and biologically meaningful cell representations.

Finally, the refined embeddings are used for clustering. By integrating the multi-omics data in a unified framework, the embeddings capture the complex relationships and interactions across different types of data, leading to more accurate and robust cell clustering results. Specifically, we employ the K-means clustering algorithm, which aims to partition the *N* cells into *M* clusters. The objective function of K-means clustering is defined as: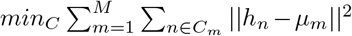, where *C*_*m*_ is the set of cells in the *m*-th cluster, is the embedding of cell *n*, and *µ*_*m*_ is the centroid of the *m*-th cluster. The centroid *µ* is calculated as: 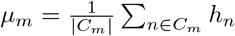.

## IV. Experiment

### A. Datasets and Evaluation Metrics

The samples used in our study come from the publicly available mouse breast cancer dataset (GSE212482) [36]. This dataset includes matched scRNA-seq, scATAC-seq, and ST data collected from the same breast tissue sections at different cancer progression stages. Specifically, tissues were dissociated into single cells using a combination of enzymatic and mechanical methods optimized to maintain cell viability while minimizing selection bias. We select subsets C3, L2 and S2, encompassing three stages : non-breast cancer, pre-breast cancer, and post-breast cancer. The details of the datasets are shown in Table I. The number of cells refers to the cells in scRNA-seq and scATAC-seq data, and the number of spots refers to the spots in ST data.

**TABLE I:** Dataset Details.

| Dataset | C3 | S2 | L2 |
| --- | --- | --- | --- |
| Initial number of genes | 83309 | 122617 | 146268 |
| Genes after preprocessing | 7044 | 49712 | 53682 |
| Number of cells | 3545 | 7616 | 5609 |
| Number of spots | 474 | 1660 | 1647 |

To evaluate clustering performance, we employ three complementary metrics: Silhouette Coefficient (SC) [30], Davies-Bouldin Index (DBI) [31], [32], and Calinski-Harabasz Index (CH) [33]. SC (ranging from -1 to 1) measures cluster cohesion and separation, with higher values indicating better clustering; DBI evaluates cluster compactness through similarity ratios, where lower values suggest better performance; and CH assesses the ratio of between-cluster to within-cluster dispersion, with higher values indicating more distinct clusters.

Modules A and C are implemented in R language, and module B is implemented in Python.

### B. Comparison on Overall Clustering Performance

Since our method is the first to integrate scRNA-seq, scATAC-seq, and ST data for clustering, the selected comparison models are those that cluster using one or two of these omics data types. Depending on the type of data used, the comparison models can be categorized into two groups: models based on scRNA-seq and scATAC-seq data: Deepmaps [27], Seurat [17], MOFA+ [13], models based on ST data: Stlern [18], CCST [28], SEDR [37].

It is important to note that the datasets used in this study are unlabeled, and all methods employed unsupervised clustering techniques. This means that there is no ground truth available to determine whether the clustering correctly identifies the corresponding cell types, so we determine the quality of clustering by the effect of clustering (such as the closeness of the same cell type, the distance between different cell types, etc.). The metrics comparison results are summarized in Table II. Additionally, we visualize the clustering results of the original datasets, the sub-optimal model MOFA+, and MOH for comparison, as shown in Fig.2. The cell type differences observed between C3, S2, and L2 samples in Fig. 2 A-C reflect the biological progression of breast cancer rather than technical artifacts from sample preparation. For example, fibroblasts and T cells dominate in the C3 stage representing normal tissue composition, while activated fibroblasts increase in the S2 stage indicating early stromal changes. Myoepithelial cells emerge prominently in L2 samples, consistent with their documented role in creating a supportive microenvironment for advanced breast cancer. This progression of cell populations aligns with known biological changes during breast cancer development, validating both our sample preparation methods and analytical approach. The results reveal several key insights.

**TABLE II:** Comparison Experiment on Three Datasets.

|  |  | Deepmaps | Seurat | MOFA+ | Stlearn | CCST | SEDR | MOH |
| --- | --- | --- | --- | --- | --- | --- | --- | --- |
| C3 | SC | 0.09 | -0.026 | 0.238 | 0.084 | 0.192 | <u>0.298</u> | **0.315** |
|  | DBI | 2.875 | 1.749 | 1.787 | 2.217 | <u>1.558</u> | 3.343 | **1.140** |
|  | CH | 279.991 | 272.081 | <u>3784.266</u> | 152.025 | 561.184 | 1150.034 | **4005.527** |
| S2 | SC | 0.026 | 0.049 | 0.062 | 0.095 | 0.049 | **0.239** | 0.236 |
|  | DBI | 7.284 | <u>2.563</u> | 3.153 | 3.528 | 3.832 | <u>2.842</u> | **2.167** |
|  | CH | 140.001 | 770.041 | <u>1359.384</u> | 169.568 | 291.331 | 1160.316 | **1364.273** |
| L2 | SC | 0.09 | -0.044 | 0.086 | 0.087 | 0.047 | <u>0.296</u> | **0.350** |
|  | DBI | 3.089 | <u>2.299</u> | 2.874 | 3.325 | 2.755 | 2.577 | **2.019** |
|  | CH | 573.294 | 340.528 | <u>1835.250</u> | 55.087 | 612.817 | 1704.901 | **2169.224** |

Note: Bold font indicates best performance, underlined values indicate second-best performance.

**Fig. 2:**
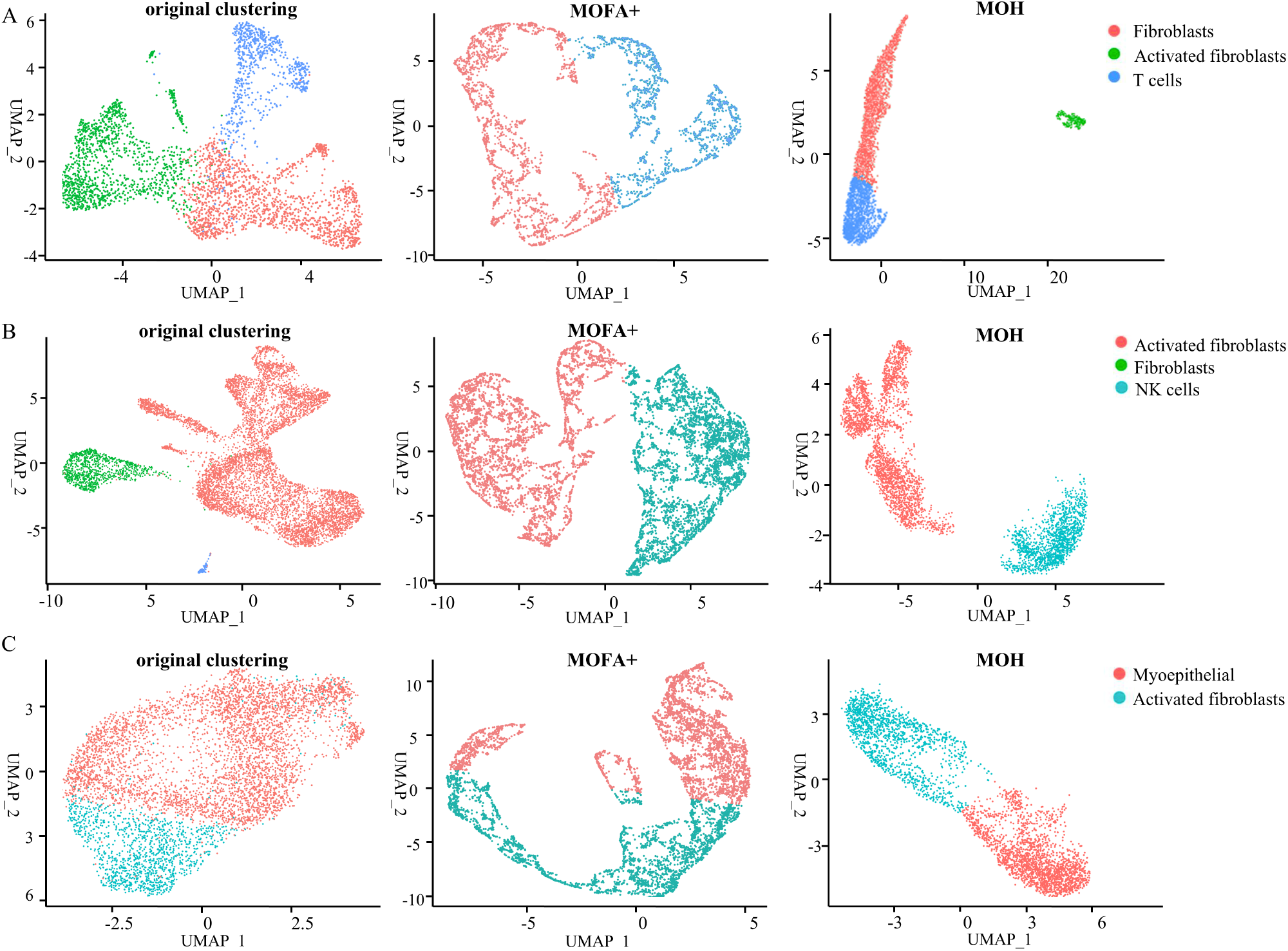
Cluster Comparison Visualization: Each color represents a different cell type. Panels A, B, and C correspond to datasets C3, S2, and L2 respectively. Columns illustrate the clustering results for different datasets using three methods: direct clustering of original data, MOFA+, and our method. The original clustering refers to direct clustering of the raw data using standard k-means clustering before any integration.

- MOH consistently achieves advanced clustering performance. For the SC metric, it outperforms the sub-optimal model by 5.70% and 18.24% on C3 and L2, respectively, and is only 1.3% below the best result on S2. This indicates that MOH forms more cohesive clusters that better reflect the underlying biological structures. For the DBI metric, which favors models with lower values indicating tighter clusters, MOH achieves reductions of 26.83%, 15.45%, and 12.18% on C3, S2, and L2, respectively. The CH metric, which measures the ratio of the sum of between-cluster dispersion to within-cluster dispersion, shows improvements of 5.85% on C3, 0.36% on S2, and 18.20% on L2. These findings underscore MOH’s robustness and consistency across different datasets, indicating its superior ability to delineate distinct clusters with tight intra-cluster cohesion and larger inter-cluster separation.
- MOH’s ability to clearly define clustering boundaries is particularly noteworthy. As illustrated in Fig. 2, MOH distinctly separates clusters, enhancing both the tightness and distinction of the clusters. This is particularly evident when comparing the clustering visualizations of the original datasets and the sub-optimal comparison model MOFA+. The visualizations highlight that MOH can delineate cluster boundaries more clearly, which is crucial for identifying subtle differences in cellular populations and understanding the underlying biological processes.
- MOH is a major advancement in multi-combination integration methods. While models like Deepmaps, Seurat, and MOFA+ focus on scRNA-seq and scATAC-seq data, and models like Stlearn, CCST, and SEDR focus on ST data, MOH consistently outperforms these models across all metrics. For instance, on C3 dataset, MOH achieves the highest SC and CH scores, and the lowest DBI score, indicating superior clustering performance. This demonstrates MOH’s superior capacity to integrate multi-omics data effectively, surpassing the performance of models limited to fewer data types.

### C. Ablation Study

Our proposed MOH model incorporates two significant innovations: 1)Defining cell-cell edges in the heterogeneous graph of scRNA-seq and scATAC-seq data. 2)Integrating ST data with scRNA-seq and scATAC-seq data to construct a multi-layer heterogeneous graph. To evaluate the effectiveness of these innovations, we conduct ablation studies defining these innovations as SMC and MLNC. We design four experimental setups: 1)Excluding both SMC and MLNC. 2)Including only SMC. 3)Including only MLNC. 4)Including both SMC and MLNC. According to TableIII, we derive the following findings.

**TABLE III:**
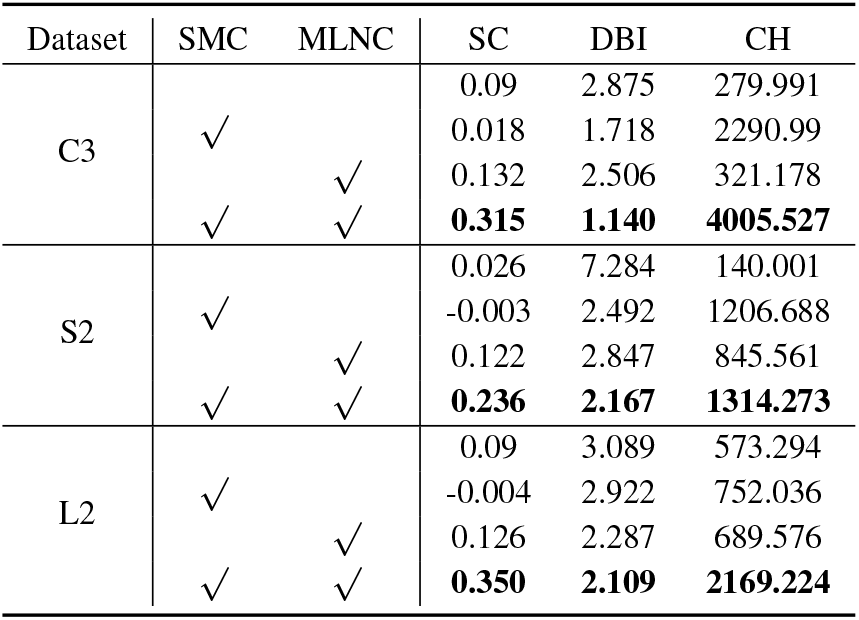
Ablation Study on Three Datasets.

- Effectiveness of adding cell-cell edges in the heterogeneous graph. Incorporating cell-cell edges in the heterogeneous graph significantly improve clustering performance across all three datasets, particularly evident in the CH metric. The process of adding cell-cell edges can be viewed as a reinforcement learning process, where the model gains more information about cell similarities through these edges, allowing it to more accurately identify and separate different cell populations. This enhanced representation capability makes the separation between clusters clearer and the cohesion within clusters tighter. Compared to the setup without cell-cell edges, the CH metric improved by 718.24%, 761.914%, and 31.18% for datasets C3, S2, and L2, respectively. Adding cell-cell edges allows structurally similar cells to enhance their embedding, resulting in more detailed comprehensive understanding of intra-cellular dynamics and interactions.
- Effectiveness of constructing a multilayer heterogeneous graph. Integrating three types of omics data to construct a multilayer heterogeneous graph result in substantial improvements across all three evaluation metrics on all datasets. It demonstrates the importance of incorporating ST-omics data. For instance, on S2 dataset, compared to using a single-layer heterogeneous graph with only two types of omics data, our method is improved by 369.23%, 70.25%, and 503.97% on SC, DBI, and CH metrics, respectively. The comprehensive multilayer heterogeneous structure takes more enriched omics data providing the comprehensive node information. The establishing interlayer connections extend the spatial context to the original data and enhance the learning of structural information, node features, and spatial relationships. The seamless integration preserves the unique characteristics of each omics data type.

### D. Parameter Sensitivity Analysis

To evaluate the sensitivity and optimal settings of MOH’s key parameters, we conducted extensive experiments focusing on the cross-layer similarity learning rate *β*, which controls the contribution of inter-layer learning in our multi-layer heterogeneous graph. We tested *β* values ranging from 0.1 to 0.9 with a step size of 0.1, while keeping other parameters fixed.

Fig 3 shows the performance metrics (SC, DBI, and CH) across different *β* values on all three datasets (C3, S2, and L2). The results demonstrate that *β* = 0.4 consistently achieves optimal performance across all evaluation metrics and datasets. This suggests that a moderate level of cross-layer interaction (40% contribution) provides the best balance for integrating information across different omics layers.

**Fig. 3:**
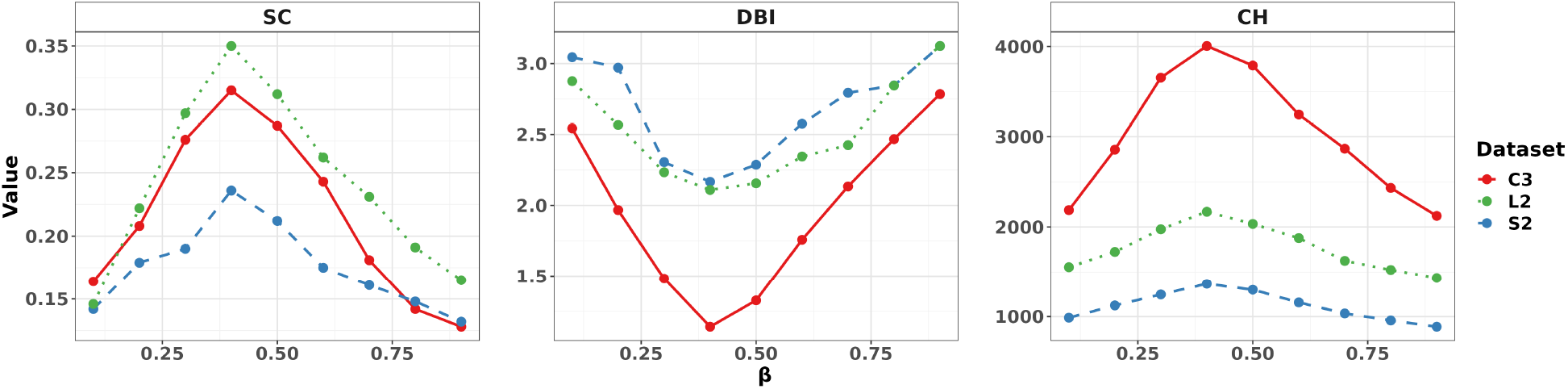
Parameter sensitivity analysis of the cross-layer similarity learning rate (*β*) on three datasets. The performance is evaluated using three metrics: Calinski-Harabasz Index (CH), Davies-Bouldin Index (DBI), and Silhouette Coefficient (SC).

### E. Gene Differential Expression Analysis

We validate MOH’s clustering accuracy and demonstrate its downstream applications. Using the Seurat package [17], we perform differential gene expression analysis to identify cell type-specific feature genes. We validate these feature genes against published literature. In the following section, we will conduct gene set enrichment analysis to uncover commonly enriched terms among these differentially expressed genes, validate them with published literature, and generate novel hypotheses for further investigation.

First, we use the FindAllMarkers function to identify all potential marker genes expressed in more than 10% of cells. Then, we rank these genes based on their differential expression values, selecting those with |*logFC*| *>* 0.5 and *p <* 0.01 as marker genes. The number of marker genes in each cell type is shown in Table IV. We choose the top three genes with the highest differential values of each cell type for visualization, with overlaps between different cell types. We visualize their expression levels within different cell types in violin plots and analyze the distribution of each marker gene on each dataset (as shown in Fig. 4).

**TABLE IV:** Details of Marker Genes on Three Datasets.

| Dataset | Celltype | Count |
| --- | --- | --- |
| C3 | Fibroblasts | 268 |
|  | Activated fibroblasts | 273 |
|  | T cells | 125 |
| S2 | Activated fibroblasts | 706 |
|  | NK cells | 717 |
| L2 | Myoepithelial | 1318 |
|  | Activated fibroblasts | 1785 |

**Fig. 4:**
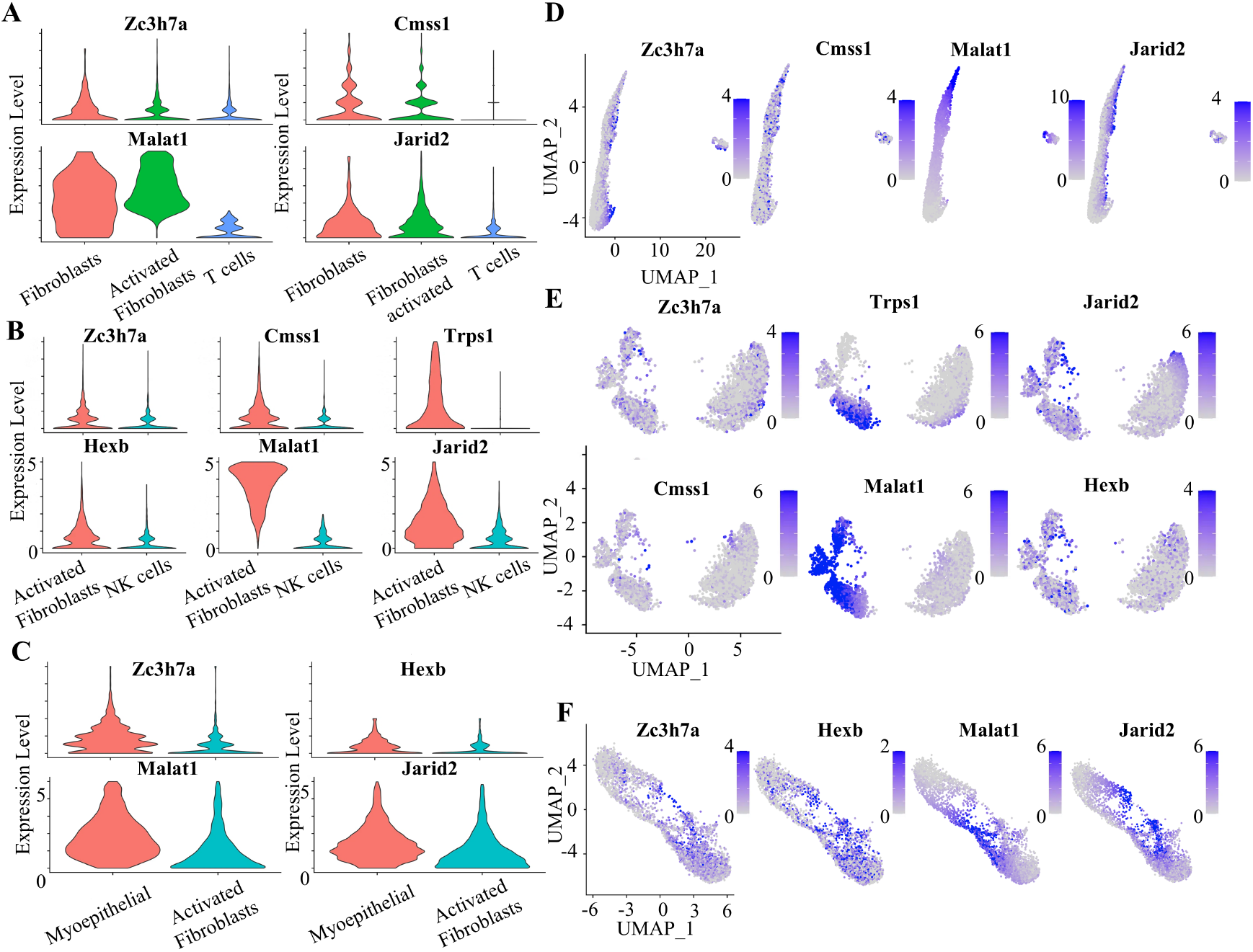
The violin plots of marker genes for datasets C3, S2, and L2 are shown in subfigures A, B, and C, displaying the expression probability density in each cluster. Their corresponding feature distribution plots are shown in subfigures D, E, and F, where color intensity in the UMAP embedding indicates expression levels, with darker shades representing higher expression.

The C3 dataset (mouse non-tumor tissue) reveals three cell types with distinct marker gene expression patterns (Fig.4A, D), including Zc3h7a, Cmss1, Malat1, and Jarid2.Cmss1 and Jarid2 also show differential expression across cell types, with Cmss1 being notably lower in T cells, further emphasizing the unique functions of each cell type. Through this analysis, we effectively identify and characterize these cell types, suggesting distinct biological functions and regulatory mechanisms within each, underscoring the biological heterogeneity of the C3 dataset.

The S2 dataset, representing precancerous breast tissue in mouse, reveals two cell types. We analyze the differential expression of six marker genes. The violin plots in Fig.4B and the distribution in Fig.4E show how these genes vary across cell types. The marker genes show distinct expression profiles between activated fibroblasts and NK cells, demonstrating the effectiveness of our clustering approach in identifying celltype specific markers.

The L2 dataset analysis reveals interesting patterns in gene expression between different cell types. While the expression differences between cell types are less pronounced compared to other datasets, this uniformity does not diminish their functional importance. The more homogeneous expression pattern suggests a stable and well-established cellular environment, where different cell types share similar molecular signatures. This observation demonstrates our method’s ability to detect and characterize subtle variations in gene expression patterns, even in complex tissue environments where traditional clustering methods might struggle to identify distinct populations. The consistent expression of certain genes (such as Malat1 and Jarid2) across different cell types suggests their involvement in fundamental cellular processes, while the overall uniformity in expression patterns may indicate a highly integrated cellular network (see Fig.4C, Fig.4F).

Since the three datasets consist of activated fibroblasts cells at different cancer stages, we expect the differentially expressed (DE) genes commonly identified across all three datasets to be associated with features of fibroblast activation. For example, Malat1 is involved in cell proliferation and migration, making it a relevant gene to study in activated fibroblasts, particularly in pathological conditions such as cancer and fibrosis [38]. Neat1 is involved in cellular processes relevant to fibroblast activation, such as stress response and cell proliferation. Its expression can indicate cellular activation states, including those found in activated fibroblasts during tissue remodeling and repair [39]. The identification of such common DE genes is validated by independent studies, supporting the accuracy of our clustering results.

The DE genes that appear exclusively in the late-stage cancer dataset are likely related to cancer progression or the role of activated fibroblasts cells in cancer. Two examples identified in our analysis are EGFR and IGFBP5. EGFR (Epidermal Growth Factor Receptor) signaling plays a crucial role in breast cancer progression and can be involved in the activation of cancer-associated fibroblasts [40] [41]. IGFBP5 (Insulin-like Growth Factor Binding Protein 5) regulates IGF signaling, which can influence breast cancer progression and may be affected by cancer-associated fibroblasts [42].

### F. Enrichment Analysis

Instead of analyzing each individual gene in a list of differentially expressed (DE) genes, an alternative approach is enrichment analysis, which identifies enriched terms that describe common characteristics of genes within a given gene set (e.g., a list of DE genes for a specific cell type). To explore the functional significance of cell type clustering, we performed Gene Ontology (GO) [43] enrichment analysis and Kyoto Encyclopedia of Genes and Genomes (KEGG) [44] pathway analysis on the marker genes identified in each cell type annotated by MOH. Fig 5A to 5F present the enrichment analysis results for the non-tumor (C3) mouse dataset, the pre-malignant (S2) breast cancer dataset, and the late-stage (L2) breast cancer dataset. The significance of each pathway represents the statistical enrichment of differentially expressed genes (−log10 p-value), with darker colors indicating stronger enrichment of cancer-related genes. Of particular note is the MAPK pathway, which shows varying significance across stages (B, D, and F) due to its dynamic role in cancer progression - from baseline activity in normal tissue to enhanced activation in advanced cancer stages [47].

**Fig. 5:**
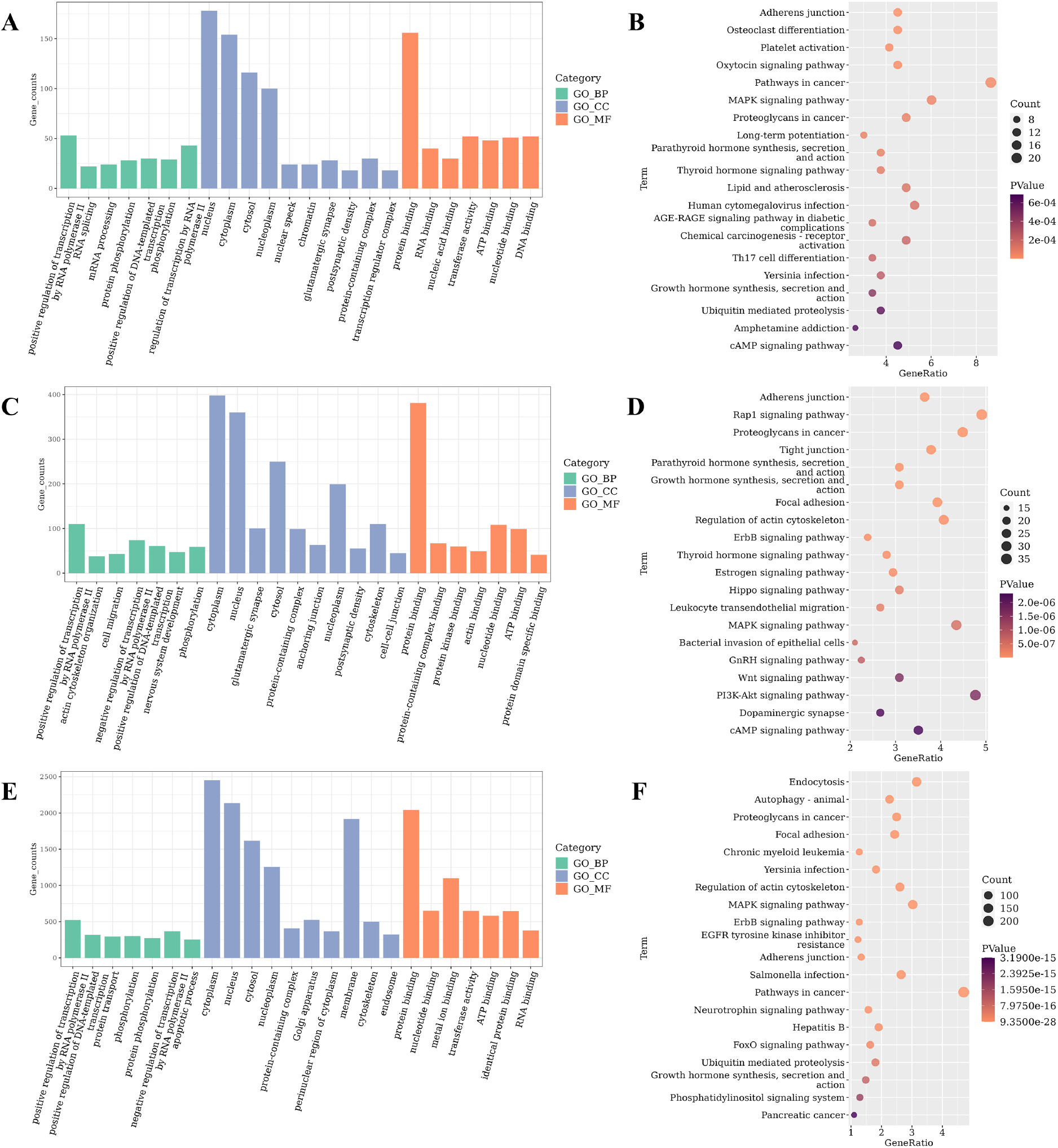
Gene Enrichment Analysis. The Gene Ontology (GO) enrichment analysis on datasets C3, S2, and L2 are respectively shown in subfigure A, C, and E. The color of the column bar represents the different filling colors of the three GO categories (BP, CC, MF). The KEGG pathway analysis on datasets C3, S2, and L2 are respectively shown in subfigure B, D, F, with the enrichment analysis results of the top 20 KEGG pathways. The bubbles represent the number of genes associated with each pathway, with larger bubbles indicating a greater number of genes. The color intensity of the bubbles indicates the pathway’s statistical significance, with darker colors representing higher significance.

In the enrichment analysis, we identified six terms that are enriched across all three stages, supporting their association with activated fibroblasts based on published studies: (1) Proteoglycans in Cancer (mmu05205), which are involved in cancer-related pathways that reflect dynamic changes in the tumor microenvironment, where fibroblasts play a supportive role in cancer progression [45]. (2) Positive Regulation of Transcription by RNA Polymerase II (GO:0045944), associated with the upregulation of gene expression, a critical process during cell activation and response to stimuli [46]. (3) Positive Regulation of DNA-templated Transcription (GO:0045893), related to the enhancement of transcription processes essential for cell activation and differentiation [46]. (4) Adherens Junction (mmu04520), which is linked to fibroblast activation through changes in cell morphology and interactions with the extracellular matrix, crucial for tissue remodeling and repair [45]. (5) Phosphorylation (GO:0016310), a key post-translational modification that regulates protein activity and signaling pathways [47]. (6) MAPK Signaling Pathway (mmu04010), involved in cell proliferation, differentiation, and stress responses, frequently enriched in activated fibroblasts due to its role in mediating responses to external signals and promoting cellular changes [47].

The terms that are only enriched in the late stage are likely related to cancer progression or the role of activated fibroblasts in cancer. This aspect of downstream analysis is particularly important, as it has the potential to generate novel hypotheses for further investigation. Our enrichment analysis identified 10 such terms, including Chronic Myeloid Leukemia (mmu05220), Yersinia Infection (mmu05135), EGFR Tyrosine Kinase Inhibitor Resistance (mmu01521), Salmonella Infection (mmu05132), Neurotrophin Signaling Pathway (mmu04722), Hepatitis B (mmu05161), FoxO Signaling Pathway (mmu04068), Phosphatidylinositol Signaling System (mmu04070), Pancreatic Cancer (mmu05212), and Protein Transport (GO:0015031). We categorize and describe these terms below, with literature support.

Role of activated fibroblasts in cancer: The term EGFR Tyrosine Kinase Inhibitor Resistance (mmu01521) is particularly relevant to activated fibroblasts in late-stage breast cancer. Cancer-associated fibroblasts (CAFs) play a crucial role in the tumor microenvironment and can contribute to drug resistance [48]. The MEK/ERK signaling pathway, involved in EGFR tyrosine kinase inhibitor resistance, has been shown to be important in the generation of CAFs [48].

Complications of immune system disorders in Cancer: Three terms are related to infections, including Yersinia Infection (mmu05135), Salmonella Infection (mmu05132), and Hepatitis B (mmu05161). This indicates a high risk of complications of immune system disorders in Cancer patients, which is confirmed by an independent study showing CAFs can create an immunosuppressive environment within the tumor microenvironment [49].

Shared characteristics of cancers: The enrichment of Chronic Myeloid Leukemia (mmu05220) and Pancreatic Cancer (mmu05212) suggests that breast cancer shares certain CAF-related characteristics with these cancers. This highlights the potential to repurpose CAF-related treatments between these cancer types, offering new avenues for therapeutic strategies [50] [51] [52]. Novel signalling pathway of cancer: There is no existing literature linking the Neurotrophin Signaling Pathway (mmu04722) and FoxO Signaling Pathway (mmu04068) to activated fibroblasts or breast cancer. Therefore, these pathways represent potential novel areas for further investigation and validation in future studies.

In summary, the enrichment results validate our cell type annotations and demonstrate the utility of MOH in identifying biologically meaningful patterns. The analysis reveals both common cellular mechanisms and condition-specific pathways, providing valuable insights for future research directions.

### G. Case Study: Breast Cancer Progression Analysis

Using our marker gene and enrichment analysis results, we present a comprehensive case study of breast cancer progression across different stages. Our datasets (C3, S2, and L2) represent a natural progression from normal tissue through precancerous to late-stage breast cancer, providing valuable insights into disease development.

Marker gene expression patterns: The progression of breast cancer shows distinct patterns in gene expression. In normal tissue (C3), we observe clear differentiation in gene expression between cell types, as shown in Fig. 4A and D, particularly in Malat1 expression between fibroblasts and T cells. As the tissue transitions to precancerous state (S2), specific genes like Trps1 show elevated expression in activated fibroblasts (Fig. 4B, E), suggesting early tumorigenic changes. In late-stage cancer (L2), we observe a more uniform expression pattern across cell types (Fig. 4C, F), indicating a mature tumor microenvironment.

Stage-specific molecular signatures: Our enrichment analysis (Fig. 5) reveals stage-specific molecular signatures. The MAPK signaling pathway shows varying significance across stages (Fig. 5B, D, F), reflecting its dynamic role in cancer progression. The EGFR Tyrosine Kinase Inhibitor Resistance pathway (mmu01521), exclusively enriched in late-stage cancer (Fig. 5F), suggests the development of drug resistance mechanisms. This is particularly relevant as cancer-associated fibroblasts (CAFs) contribute significantly to drug resistance through the MEK/ERK signaling pathway [48].

Therapeutic implications: The identification of shared characteristics between breast cancer and other cancer types (e.g., pancreatic cancer and chronic myeloid leukemia) suggests opportunities for drug repurposing, as revealed in our late-stage enrichment analysis (Fig. 5F),. For instance, while Galunisertib and Sunitinib are used in both breast and pancreatic cancers [50], their potential application in chronic myeloid leukemia warrants investigation. Similarly, Plerixafor, currently used only in breast cancer [51], might be effective in treating other cancer types.

Novel pathways: Our analysis identified previously unexplored pathways in breast cancer progression, particularly the Neurotrophin and FoxO signaling pathways(Fig. 5F). These findings suggest new directions for investigating breast cancer mechanisms and potential therapeutic targets.

## V. Discussion

MOH introduces a pioneering approach for single cell clustering by establishing a novel multilayer heterogeneous graph architecture that uniquely integrates three multi-omics data, including scRNA-seq, scATAC-seq, and ST data. This innovation addresses key challenges in multi-omics integration while demonstrating superior performance in both technical implementation and biological insights generation.

### A. Advantages

Superior integration performance: MOH demonstrates exceptional performance in integrating heterogeneous omics data, as validated by our comparative experiments against six state-of-the-art methods. The superior clustering performance on three real world datasets particularly highlights MOH’s effectiveness in preserving both spatial and molecular information during integration.

Enhanced feature learning through graph structure: Our ablation studies and enrichment analysis demonstrate that incorporating additional cell-cell edges significantly enhances the model’s capabilities in two aspects: (1) improved clustering performance compared to traditional heterogeneous graphs with only cell-gene edges, and (2) enhanced molecular pattern recognition, as evidenced by our ability to detect dynamic changes in MAPK signaling pathway activity across different cancer stages. These improvements directly result from the model’s enhanced ability to capture and propagate cellular interaction information through the enriched graph structure.

Domain adaptation and transfer learning capabilities: MOH exhibits effective domain adaptation through its graph-based knowledge transfer between omics layers and self-supervised learning approach. Its consistent performance across multiple datasets demonstrates robust generalization to varied experimental conditions, while the flexible architecture supports transfer learning by enabling seamless integration of new omics data without structural redesign.

Biological discovery capability: MOH has significant clinical potential in cancer subtype identification, therapeutic target discovery, disease monitoring, and drug development.The enrichment analyse and case study validate MOH’s biological relevance through the identification of 10 uniquely enriched terms in late-stage cancer. These findings led to novel insights into immune system complications in cancer and the discovery of previously unexplored signaling pathways in breast cancer.

### B. Limitations and Future Work

MOH’s flexible architecture enables seamless integration of additional omics layers, allowing adaptation to new datasets or experimental conditions with minimal modification. This adaptability positions MOH as a strong candidate for domain adaptation and transfer learning, where knowledge from well-characterized datasets can be leveraged to improve performance on less-explored ones, enhancing generalization across diverse biological contexts. While experimental results demonstrate the superior performance of MOH, its computational complexity—particularly the O(n^2^) time for cross-layer cell similarity calculation—poses a scalability challenge for increasingly large single-cell datasets. Future work will focus on optimizing this step through approximate methods to ensure scalability without compromising biological interpretability.

## VI. Conclusion

In this study, we introduce MOH, a novel multi-layer, multiomics heterogeneous graph framework designed for single-cell clustering. Different from previous works, it integrates more than two omics. MOH offers a scalable approach for integrating multiple omics types into a unified framework, enabling more accurate cross-omics learning. The comparative experimental results demonstrate that MOH’s multi-layer structure and inter-layer node communication improve the clustering performance, providing a more comprehensive analysis of cellular features. Our comprehensive ablation studies enhance the effectiveness of both cell-cell edges and multilayer structure design. Through application to breast cancer progression analysis, we revealed stage-specific molecular signatures and identified novel therapeutic targets, particularly in the EGFR Tyrosine Kinase Inhibitor Resistance pathway, offering new insights into cancer treatment strategies. The superior performance across multiple datasets demonstrates MOH’s potential for advancing single-cell multi-omics research and its clinical applications.

## AVAILABLE

The codes and supplementary document are available at https://github.com/hebutdy/MOH.

## Notes

This work was supported in part by Natural Science Foundation of Tianjin(23JCYBJC00790), Chinese State Scholarship Fund(202108130108)

### Competing Interest Statement

The authors have declared no competing interest.

